# Drug susceptibility testing of *Mycobacterium simiae*: An emerging pathogen in Iran

**DOI:** 10.1101/549428

**Authors:** Mohammad Javad Nasiri, Sirus Amini, Zahra Nikpor, Samaneh Arefzadeh, Seyed Mohammad Javad Mousavi

**Author notes:** Corresponding author: Department of Microbiology, School of Medicine, Shahid Beheshti University of Medical Sciences, Tehran, Iran. Author’s contribution: MJN: Designed the protocol MJN, SA, ZN, SA: Performed the laboratory procedures and statistical analysis MJN, SMJM: Wrote the first draft of the manuscript MJN: Revised the manuscript.

## Abstract

**Introduction:** *Mycobacterium simiae* is an emerging pathogen in Iran and little is known about drug susceptibility patterns of this pathogen.

**Materials and methods:** Twenty five clinical isolates of *M. simiae* from 80 patients with confirmed NTM pulmonary disease were included in this study. For drug susceptibility testing (DST), proportional and broth microdilution methods were used according to the clinical and laboratory standards institute (CLSI) guideline.

**Results:** All clinical isolates of *M. simiae* were resistant to isoniazid, rifampicin, ethambutol, streptomycin, amikacin, kanamycin, ciprofloxacin and clarithromycin. They also were highly resistant to ofloxacin (80%). Susceptibility to ofloxacin was only noted in the 5 isolates.

**Conclusions:** Clinical isolates of *M*. *simiae* were multidrug resistant, and had different drug susceptibility patterns than previously published studies. DST results can assist in selecting more appropriate treatment regimens. Newer drugs with proven clinical efficacy correlating with in vitro susceptibility should be substituted with first- and second line anti-TB drug testing.

**Author summary:** *Mycobacterium simiae* can cause serious pulmonary diseases. It has been recognized as an emerging pathogen in Iran. Although few cases have been reported, *M. simiae* has been associated with multidrug-resistance. In our study, we have observed a surprisingly high proportion of highly drug-resistant *M. simiae*. This data call for a scale-up of TB control program in Iran.

## Introduction

Infections caused by nontuberculous mycobacteria (NTM) have been recently reported as an important public health problem in many parts of the world, especially in developing countries [1]. In Iran, a country with a moderate rate of tuberculosis (TB), *Mycobacterium fortuitum*, *M. kansasii*, and *M. simiae* are the most common causes of NTM diseases, respectively [2, 3]. *M. simiae* is becoming an emerging pathogen in Iran and has been recognized as a causative agent of pulmonary and disseminated infections [4, 5]. Treatment of *M. simiae* disease is time-consuming and often complicated [6, 7]. Based on previous studies, *M. simiae* exhibited the highest level of drug resistance to several anti-TB agents in vitro [7, 8]. Although the association between in vitro susceptibility and the in vivo treatment outcome has not been established for most drugs, treatment of individual patients should preferably be based on drug susceptibility testing (DST) results [7]. Given the limited data on drug susceptibility patterns of clinical *M. simiae* isolates in Iran, we aimed to report the DST results of *M. simiae* isolated from clinical samples and discuss the implications of these findings for treatment regimens.

## Materials and Methods

### Samples and setting

Twenty-five clinical isolates of *M. simiae* were included in this study. These samples were recovered from 80 patients with NTM pulmonary diseases who referred to Tehran regional TB reference laboratory (TRTB-RL) from March 2014 to January 2018. The remaining 55 species were other NTM. The mean age of patients was 51.6 ± 19.3 years. *M*. *simiae* was isolated from respiratory specimens (sputum, bronchoalveolar lavage). All clinical isolates form patients with NTM met ATS/IDSA diagnostic criteria for inclusion and were considered as the causative pathogen for the disease [9]. TRTB-RL is well-equipped with biosafety level III facilities and is able to perform DST.

### Phenotypic and molecular Identification of M. simiae

Growth rate, macroscopic and microscopic morphological features, growth at different temperatures and also a set of biochemical tests including tween 80 hydrolysis, nitrate reduction, niacin production, arylsulfatase, urease production, tellurite reduction, salt tolerance, and catalase production were used for identification of *M. simiae* [10].

For molecular identification, PCR restriction analysis (PRA) and sequencing of 16S rRNA were used as described previously [11, 12].

### DST

Two methods were used for DST; proportional assay and broth microdilution method. The following antibiotics were selected: isoniazid, rifampicin, ethambutol, streptomycin, amikacin, kanamycin, ciprofloxacin, ofloxacin, and clarithromycin were selected. The stock solutions of each agent were prepared by dissolving pure powder of individual drugs in sterile distilled water or another suitable solvent.

### Proportional DST

In proportional DST, resistance was expressed as the percentage of colonies that grew on critical concentrations of the drugs according to the guidelines established by the clinical and laboratory standards institute (CLSI) [13]. The interpretation was made according to the usual criteria for resistance, i.e., 1% for all drugs. *M. tuberculosis* H37Rv strain (ATCC 27294) was used for quality control testing in DST [14].

### Broth microdilution

For each isolate, a suspension in the broth was prepared from an agar medium with adequate growth of the isolate and the inoculum was standardized for susceptibility testing to a turbidity equivalent to a 0.5 McFarland standard. Suspensions were then diluted and inoculated into 96-well microtiter plates to achieve a final organism concentration from 1 × 10^5^ to 5 × 10^5^ CFU/ml. The antimicrobial agent minimum inhibitory concentrations (MICs) were interpreted according to the guidelines established by the CLSI [13]. Serial double dilutions of antimicrobial agents were prepared ranging from 0.06 to 512 mg/L, added to cation-supplemented Mueller–Hinton broth plus 5% OADC, and then 0.1 ml was dispensed into the wells of microdilution plates. Following inoculation, all cultures were incubated aerobically at 37 °C. Growth was examined weekly up to 4 weeks. The MIC was defined as the lowest concentration of the drug that inhibited visible growth. Quality control of MICs was performed by testing recommended reference strains, including *Enterococcus, faecalis* ATCC 29212, *Mycobacterium peregrinum* ATCC 700686, *Pseudomonas aeruginosa* ATCC 27853, and *Staphylococcus aureus* ATCC 29213.

### Data storage and analysis

Information and laboratory data were entered in Microsoft Excel 2017 software and were imported into SPSS (version 22; IBM) software. Frequency distributions were analyzed by the use of the SPSS (version 22) software.

## Ethics

All adult subjects provided informed written consent, and a parent or guardian of any child participant provided informed consent on the child’s behalf. The Ethics Committee of Shahid Beheshti University of Medical Sciences approved the study (approval number: IR.SBMU.RETECH.REC.1397.245).

## Results

Based on the results of proportional DST, all 25 (100%) isolates of *M. simiae* were resistant to isoniazid, rifampicin, ethambutol, streptomycin, amikacin, kanamycin, ciprofloxacin, and clarithromycin (Table 1 and 2). They also were highly resistant to ofloxacin (80%). Susceptibility to ofloxacin was only noted in the 5 isolates. In vitro susceptibility to first-line anti-TB drugs was not found among clinical isolates of *M. simiae*. Acquired resistance, defined as a resistant follow-up isolate after a susceptible primary isolate, was noted in 10 patients; acquired rifampicin resistance was most frequent, followed by acquired and isoniazid resistance.

**Table 1.** In vitro resistance of the 25 *M. simiae* isolates to the tested antibiotics by proportional method.

| Antibiotic | No. (%) of isolates |  |  |
| --- | --- | --- | --- |
|  | Susceptible | Intermediate | Resistant |
| Isoniazid | 0 (0) | - | 25 (100) |
| Rifampicin | 0 (0) | - | 25 (100) |
| Ethambutol | 0 (0) | - | 25 (100) |
| Streptomycin | 0 (0) | - | 25 (100) |
| Amikacin | 0 (0) | - | 25 (100) |
| Kanamycin | 0 (0) | - | 25 (100) |
| Ciprofloxacin | 0 (0) | - | 25 (100) |
| Ofloxacin | 5 (20) | - | 20 (80) |
| Clarithromycin | 0 (0) | - | 25 (100) |

**Table 2.** In vitro resistance of the 25 *M. simiae* isolates to the tested antibiotics by broth microdilution method.

| Antibiotic | Range | MIC (lg/ml) |  | No. (%) of isolates |  |  |
| --- | --- | --- | --- | --- | --- | --- |
|  |  | MIC50 | MIC90 | Susceptible | Intermediate | Resistant |
| Isoniazid | 16–128 | 32 | 64 | 0 (0) | - | 25 (100) |
| Rifampicin | 1-128 | 4 | 64 | 0 (0) | - | 25 (100) |
| Ethambutol | 4-64 | 16 | 32 | 0 (0) | - | 25 (100) |
| Streptomycin | 2-64 | 16 | 32 | 0 (0) | - | 25 (100) |
| Amikacin | 0.25-64 | 16 | 32 | 0 (0) | - | 25 (100) |
| Kanamycin | 0.25-64 | 16 | 32 | 0 (0) | - | 25 (100) |
| Ciprofloxacin | 0.25-64 | 8 | 32 | 0 (0) | - | 25 (100) |
| Ofloxacin | 1-64 | 8 | 32 | 5 (20) | - | 20 (80) |
| Clarithromycin | 32–128 | 32 | 64 | 0 (0) | - | 25 (100) |

## Discussion

*M. simiae* as an emerging pathogen in Iran accounts for more than 30.0% of all pulmonary diseases caused by NTM in our study. Based on the previous studies, 10 to 47% of the patients with NTM had true *M*. *simiae* infection [5, 15, 16]. We have recently indicated the clinical isolation of *M*. *simiae* by some studies in Iran, but the ecology behind this is not well understood [17]. We hypothesize that the high rate of *M. simiae* in Iran may be due to the contamination of water supplies with this pathogen, a finding that was previously reported in studies conducted in Iran [18, 19]. Temperature and humidity have also been reported to be associated with NTM infections [16, 20]. Another explanatory factor may be the presence of underlying diseases among patients with *M. simiae* as described by Coolen-Allou [20].

Another issue raised by *M. simiae* pulmonary disease is its difficulty in treatment. Since little is known about drug susceptibility patterns of clinical isolates of *M. simiae* in Iran, there is little basis to predict potentially successful treatment regimens. Based on our study, all *M. simiae* species showed resistance to isoniazid, rifampicin, ethambutol, streptomycin, amikacin, kanamycin, ciprofloxacin, clarithromycin and, to lesser extent ofloxacin. Van Ingen et al in the Netherlands showed resistant rates of 86% to amikacin, 77% to ciprofloxacin, 36% to moxifloxacin and 9% to clarithromycin in clinical isolates of *M. simiae* [7]. *M. simiae* isolates from Hamieh et al, a study in Lebanon were mostly susceptible to amikacin and clarithromycin and less susceptible to moxifloxacin and ciprofloxacin [16]. Likewise, in a study conducted by Coolen-Allou et al, in the France, *M. simiae* isolates were found to have low susceptibility to antibiotics, except for amikacin, fluoroquinolones, and clarithromycin [20]. These findings indicate the variable drug susceptibility patterns of *M. simiae* in different geographic locations and emphasize on the need to perform DST before initiating therapy.

This study has some limitations. First, the small numbers of evaluated patients and events might have reduced the power of the study. Second, the potential effect of predisposing factors could not be analyzed because of the limited information obtained from our data sources. The results of this single-center study cannot be extrapolated for the whole country.

In conclusions, clinical isolates of *M*. *simiae* were multidrug resistant and had different drug susceptibility patterns than previously published studies. DST results can assist in selecting more appropriate treatment regimens. Newer drugs with proven clinical efficacy correlating with in vitro susceptibility should be substituted with first- and second line anti-TB drug testing. Further molecular studies on environmental and clinical isolates are needed to better assess the epidemiology and ecology of *M. simiae*.

## Acknowledgements

This study was supported by Shahid Beheshti University of Medical Sciences, Tehran, Iran.

